# Expansion microscopy facilitates quantitative super-resolution studies of cytoskeletal structures in kinetoplastid parasites

**DOI:** 10.1101/2021.04.20.440601

**Authors:** Peter Gorilak, Martina Pružincová, Hana Vachova, Marie Olšinová, Vladimir Varga

## Abstract

Expansion microscopy (ExM) has become a powerful super-resolution method in cell biology. It is a simple, yet robust approach, which does not require any instrumentation or reagents beyond those present in a standard microscopy facility. In this study, we used kinetoplastid parasites *Trypanosoma brucei* and *Leishmania major*, which possess a complex, yet well-defined microtubule-based cytoskeleton, to demonstrate that this method recapitulates faithfully morphology of structures as previously revealed by a combination of sophisticated electron microscopy (EM) approaches. Importantly, we also show that due to rapidness of image acquisition and 3D reconstruction of cellular volumes ExM is capable of complementing EM approaches by providing more quantitative data. This is demonstrated on examples of less well-appreciated microtubule structures, such as the neck microtubule of *T. brucei* or the pocket, cytosolic, and multivesicular tubule-associated microtubules of *L. major*. We further demonstrate that ExM enables identifying cell types rare in a population, such as cells in mitosis and cytokinesis. 3D reconstruction of an entire volume of these cells provided details on morphology of the mitotic spindle and the cleavage furrow. Finally, we show that established antibody markers of major cytoskeletal structures function well in ExM, which together with the ability to visualize proteins tagged with small epitope tags will facilitate studies of the kinetoplastid cytoskeleton.

## Introduction

Kinetoplastida are a group of protists, comprising both free-living and parasitic organisms. Of these, the most notorious are *Trypanosoma brucei*, the causative agent of African sleeping sickness in humans and nagana in cattle, *T. cruzi*, causing Chagas disease in Americas, and numerous species of *Leishmania* genus, causing cutaneous, mucocutaneous, and visceral leishmaniosis on several continents. These important parasites are transmitted between mammalian hosts by insect vectors. Progression of infection in both the host and the vector is dependent on the ability of the parasites to colonize, in a defined sequence, various tissues. To achieve so, transitions between various proliferative life cycle forms occur via differentiation events associated with dramatic changes to cell shape, size, and arrangement of organelles (Matthews, 2005).

Despite the considerable morphological diversity, kinetoplastid cells share a common design principle with their morphology being determined by the microtubule-based cytoskeleton; the microtubules form the subpellicular corset, an array of parallel evenly-spaced microtubules, which subtend the entire cytoplasmic membrane (Gull, 1999). Another prominent feature is the presence of the flagellum, the principal motility organelle, which contains the evolutionary conserved microtubule-based axoneme, and the kinetoplastid-specific paraflagellar rod (Gull, 1999). The flagellum specifies, via association with its base, the position of single copy organelles, such as the kinetoplast, i. e. mitochondrial DNA, and the flagellar pocket (Hayes et al., 2014). Moreover, in *T. brucei* the attachment of the flagellum to the cell body via the structure called the flagellum attachment zone (FAZ) defines cell shape, polarity, and the starting point of cytokinesis (Kohl et al., 2003). Replication of these cytoskeletal structures is coordinated, tightly linked to the cell cycle, and occurs with high fidelity (Sherwin and Gull, 1989). Thus, kinetoplastid cells of a proliferative population show a remarkable uniformity of their cytoskeletal architecture.

Studies of the kinetoplastid cytoskeleton have been heavily reliant on electron microscopy (EM), mainly due to the spacing of various structural components, such as the corset microtubules, axonemal microtubule doublets, domains of the FAZ etc., being below the resolution limit of conventional light microscopy, which is about 180 nm in the lateral and 500 nm in the axial direction (Tam and Merino, 2015). Indeed, EM provided a considerable understanding of the kinetoplastid cytoskeletal architecture (e.g. Lacomble et al., 2009; Sherwin and Gull, 1989; Wheeler et al., 2016). However, the EM approaches require specialized reagents, instrumentation, and are time demanding, making certain types of studies challenging. These include for example an ultrastructural investigation of cells which are rare in a population; yet, these may be key for understanding life cycle transition processes. Another example are studies aiming to determine localization of specific molecules in the entire cellular volume. Similar limitations also apply to major super-resolution optical microscopy techniques, such as STED and STORM (Tam and Merino, 2015).

Over recent years another super-resolution method, termed Expansion microscopy (ExM), was developed (Chen et al., 2015; Gambarotto et al., 2019). It is based on embedding a specimen into a swellable polymer and its subsequent isotropic expansion. These specimens, typically expanded 4-5 times in each direction, are thereafter imaged with a confocal microscope; this produces a corresponding increase in resolution and is comparatively rapid.

ExM has been extensively employed to study mammalian cells and tissues (e.g. Ku et al., 2016; Tillberg et al., 2016; Zhao et al., 2017). In contrast, only a limited number of studies of parasitic protists have been conducted so far. Yet, investigations of *Giardia intestinalis* (Halpern et al., 2017), *Plasmodium* (Bertiaux et al., 2021), as well as the recent study of a mitochondrial genome segregation factor in *T. brucei* (Amodeo et al., 2021), demonstrated that due to small sizes of their cells ExM offers clear opportunities for analysing structural features of these important parasites. Moreover, due to a well-defined morphology of their cellular cytoskeleton protists enable assessing the ExM approach at cellular dimensions.

In this work, we set out to optimize and validate ExM for *Trypanosoma brucei* and *Leishmania major* cells, which possess a complex, highly defined, and a relatively well-characterized microtubule-based cytoskeleton, as rehearsed above. We show that data obtained by ExM recapitulate the previous EM observations with regard to the presence, positioning, orientation, and shape of individual structures. However, caution should be observed when using ExM to determine distances, as individual structures may expand to a different extent. This was previously noticed in other experimental systems (e.g. Büttner et al., 2020; Pernal et al., 2020). Importantly, we observed that ExM enables resolving individual cytoskeletal components, such as the microtubules. This, together with the quantitative aspect of the approach facilitated better understanding of less well-appreciated structures, such as the neck microtubule *in T. brucei*, or the pocket, cytoplasmic, and multivesicular tubule-associated microtubules in *L. major*. We further observed in a subset of *L. major* cells short microtubules or plate-like structures associated with basal and pro-basal bodies. Moreover, ExM facilitated studies of rare cell types in a population, providing detailed information on the structure of the mitotic spindle and cleavage furrow in *L. major*.

Finally, we demonstrate that a number of commonly used antibody markers can be used to visualize the respective cytoskeletal structures by ExM as well as localize epitope-tagged proteins in the entire cellular volume. We envisage that due to its robustness, flexibility, and a rapid image acquisition ExM will become a valuable approach to study the biology of the kinetoplastid parasites.

## Results

### 1/ Optimizing ExM for kinetoplastid cells

To optimize and validate the ExM protocol for kinetoplastids we used procyclic *T. brucei brucei* cells, which naturally occur in the tsetse fly midgut. They can, however, be readily cultured in a laboratory and hence their morphology, cellular architecture, and changes to these during the cell cycle have been extensively characterized (Lacomble et al., 2009; Sherwin and Gull, 1989).

We employed a variation of ExM termed ultrastructure ExM, which is based on fixation of cells with 4% formaldehyde and 4% acrylamide, followed by gelation with 19% sodium acrylate, 10% acrylamide, and 0.1% N, N’-methylenebisacrylamide, denaturation with 200 mM sodium dodecyl sulphate at 95°C, and antibody staining (Gambarotto et al., 2019). Following a described protocol for handling free-swimming *Chlamydomonas* cells (Gambarotto et al., 2019), we adhered procyclic *T. brucei* cells to a glass coverslip and further processed for ExM. To visualize expanded microtubule-based structures to validate the procedure we tested several anti-tubulin antibodies: the polyclonal rabbit anti-alpha tubulin antibody (See Table 1 for specification), the mouse monoclonal anti-alpha tubulin antibody TAT1 (Woods et al., 1989), the mouse anti-beta tubulin antibody KMX-1 (Birkett et al., 1985), and V tubulin antibody C3B9 (Woods et al., 1989).

**Table 1-.** list of used antibodies.

| Primary antibody | Target | Source | Raised in | Dilution for ExM | Dilution for IF |
| --- | --- | --- | --- | --- | --- |
| polyclonal anti-alpha tubulin | alpha tubulin | Genetex GTX 15246 | rabbit | 1:20 | 1:10 |
| TAT1 | alpha tubulin | Woods et al., 1989 | mouse | 1:10 | 1:100 |
| KMX-1 | beta tubulin | Birkett et al., 1985 | mouse | 1:10 | 1:100 |
| C3B9 | acetylated alpha tubulin | Woods et al., 1989 | mouse | 1:10 | 1:100 |
| L3B2 | FAZ1 constituent of FAZ | Kohl et al., 1999 | mouse | 1:10 | 1:40 |
| anti-ClpGM6 (CUK1114) | ClpGM6 constituent of FAZ | Hayes et al., 2014 | rabbit | 1:200 | 1:200 |
| L8C4 | PFR2 constituent of PFR | Kohl et al., 2007 | mouse | 1:10 | 1:20 |
| anti-HA tag | HA tag | Cell Signaling Technology 3724S | rabbit | 1:500 | 1:500 |
| mAB25 | axonemal protein TbSAXO | Pradel et al., 2006 | mouse | 1:25 | 1:20 |
| anti-FTZC | flagellum transition zone component | Bringaud et al., 2000 | rabbit | 1:500 | 1:1000 |
| Did not produce any specific signal in ExM |  |  |  |  |  |
| BBA4 | basal body and pro-basal body | Woods et al., 1989 | mouse | 1:50 | 1:100 |
| mAb62 | flagella connector | Abeywickrema et al., 2019 | mouse | 1:25 | 1:500 |
| Secondary antibody | Source | Raised in | Dilution for ExM | Dilution for IF |  |
| Alexa Fluor 488 anti-Mouse IgG (H+L) | Invitrogen A11001 | goat | 1:500 | 1:1000 |  |
| Alexa Fluor 488 anti-Rabbit IgG (H+L) | Invitrogen A11008 | goat | 1:500 | 1:1000 |  |
| Alexa Fluor 555 anti-Mouse IgG (H+L) | Invitrogen A21422 | goat | 1:500 | 1:1000 |  |
| Alexa Fluor 555 anti-Rabbit IgG (H+L) | Invitrogen A21428 | goat | 1:500 | 1:1000 |  |

A 3D volume reconstructed from a series of confocal z-planes of expanded cells revealed that the anti-tubulin signal of each of the antibodies outlined the entire microtubule corset (Fig. 1A and Suppl. fig. 1). The corset retained its typical elongated shape, including being wider at the posterior cell end than at the anterior end, as demonstrated previously by light microscopy (Alizadehrad et al., 2015), as well as by EM (Sherwin and Gull, 1989). To estimate the expansion factor we turned the z-stacks into 2D images by applying maximum intensity projection and measured the length of the corsets; these were between 70 - 110 µm long. Corsets of formaldehyde-fixed non-expanded cells imaged by confocal microscopy (Fig. 1B) were 16 - 24 µm long, yielding the expansion factor of approximately 4.5. This is in a good agreement with the measured 4.7-fold physical expansion (4.58, 4.66, and 4.71-fold expansion in three experiments) of the gel in water prior to imaging.

**Figure 1.**
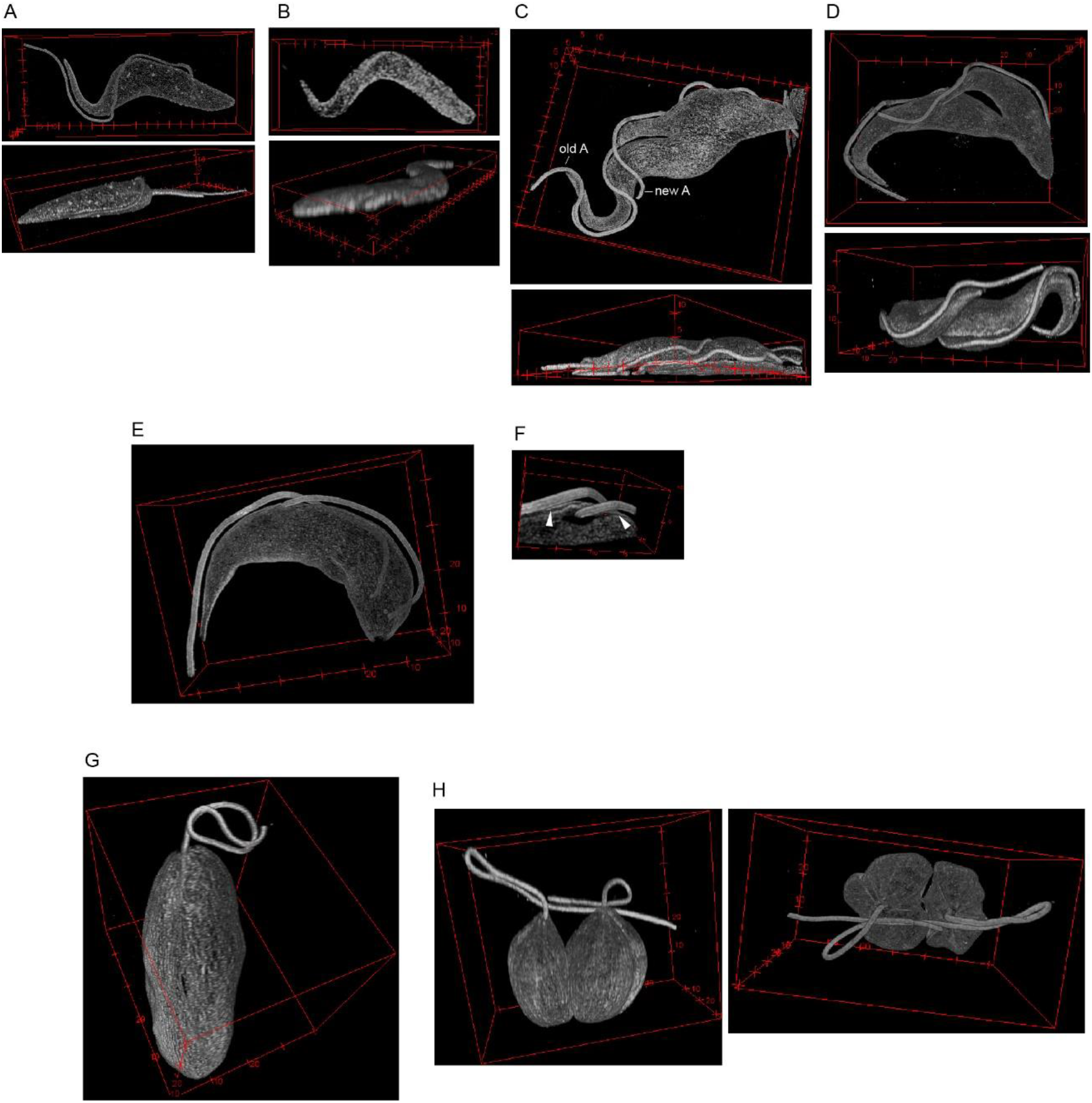
ExM visualizes 3D morphology of the subpellicular corset and the axoneme. A-3D reconstruction of a confocal z-stack of a procyclic *T. brucei* cell fixed on a coverslip and stained with the rabbit polyclonal anti-tubulin antibody. Top - en face view; Bottom - side view. Axes labels are in µms for all 3D reconstructed images. B-3D reconstruction of a confocal z-stack of a non-expanded procyclic *T. brucei* cell fixed on a coverslip and stained with the C3B9 antibody. Top - en face view; Bottom - side view. C-3D reconstruction of a confocal z-stack of a procyclic *T. brucei* cell in cytokinesis, which was fixed on a coverslip and stained with the C3B9 antibody. Top - en face view; Bottom - side view. Old A old axoneme; New A - new axoneme. D-3D reconstruction of a confocal z-stack of an expanded procyclic *T. brucei* cell in cytokinesis, which was fixed in solution before being settled on a coverslip and stained with the C3B9 antibody. Top en face view; Bottom - side view. E-En face view of 3D reconstruction of an expanded bloodstream *T. brucei* cell stained with the C3B9 antibody. F-3D reconstruction of the area around the distal end of the new flagellum. Note the groove devoid of the microtubule signal, which houses the end of the new axoneme. In addition, lines underneath both axonemes devoid of the signal (arrowheads) correspond to the position of the FAZ filament. G-En face view of 3D reconstruction of an expanded *L. major* cell with a single flagellum stained with the C3B9 antibody. H-3D reconstruction of an expanded *L. major* cell with two flagella undergoing cytokinesis. The cell was stained with the TAT1 antibody. Left - side view; Right - view from anterior end.

Tilting 3D reconstructed corsets revealed that they were significantly flattened on the side juxtaposed to the surface of the gel (Fig. 1A and 1C). We assumed this flattening resulted from the adhesion of live cells to the glass surface at the beginning of the ExM procedure. Indeed, we observed similar flattening of non-expanded cells, which were attached to glass alive (Fig. 1B). Such a reduction in the cell height may be beneficial, as imaging thick specimens is subject to a significant signal loss and aberrations (Hell et al., 1993). However, to obtain a more realistic visualization of the shape of the corset we adapted the procedure for the preparation of specimens for scanning EM (Hayes et al., 2014). Live cells were first fixed in solution, settled onto a coverslip, and thereafter processed for expansion microscopy. This indeed led to only a very limited area of a cell being flattened (Fig. 1D and Suppl. fig. 1). To mitigate the signal loss we tried to match the refractive index of the gel to the one of the oil immersion by incubating it with a solution of 84% sucrose (Gao et al., 2018). This indeed partially reduced the signal loss, but in accordance with previous observations (Gao et al., 2018) also led to a 10% reduction in the gel size with the final expansion factor of 4.3. Therefore, the approach was not universally employed in this study.

In contrast to non-expanded cells, in which the anti-tubulin antibodies stained the axonemal microtubules only weakly (Fig. 1B, Suppl. fig. 2, see also Woods et al., 1989) there was a strong signal associated with this structure in expanded specimens (Fig. 1A, 1C, 1D, and Suppl. fig. 1). Hence, the axoneme could be traced continuously from the point of its exit from the corset, running in an incomplete turn alongside the corset, and extending beyond its anterior end (Fig. 1A, 1C, 1D, and Suppl. fig. 1), consistent with previous observations (Alizadehrad et al., 2015). A gap between the corset and the axoneme was always present. This space is occupied by the cellular and flagellar membranes, and also contains the junction complex of the FAZ (Sunter and Gull, 2016) (see below for antibody staining against the FAZ constituents).

**Figure 2.**
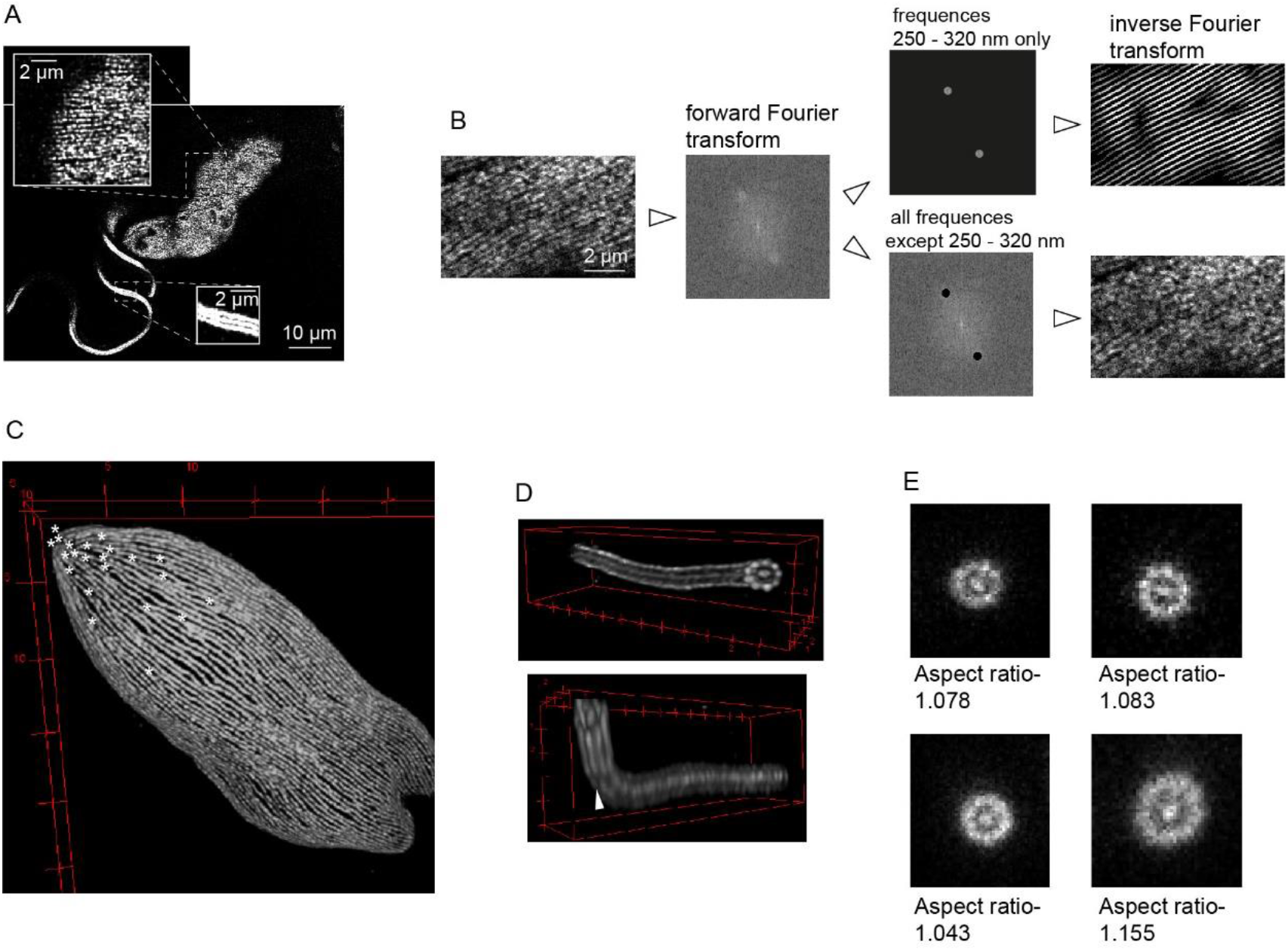
ExM resolves individual structural components. A- A single confocal plane of an expanded procyclic *T. brucei* cell fixed on a coverslip and stained with the C3B9 antibody. The insets show the microtubule corset and the axoneme, respectively. B- An example of an analysis of distances between microtubules in the subpellicular corset of procyclic *T. brucei*. Confocal z-planes encompassing corset microtubules in the vicinity of the coverslip and oriented approximately parallel to the image plane were projected to 2D by the average intensity algorithm. The forward Fourier transform of the resulting image was performed. The frequency domain image shows strongly represented frequencies with the periodicity of 250 320 nm. The inverse Fourier transform of these frequencies recapitulated the majority of the microtubule-associated signal (top), while the inverse Fourier transform of remaining frequencies did not (bottom). C- 3D reconstruction of the posterior part of the subpellicular corset of a *L. major* cell (an identical cell to Fig. 1G). Ends of individual corset microtubules are indicated with an asterisk. D- 3D reconstruction of the distal part of the axoneme of a procyclic *T. brucei* cell. Top - view from the very end with 9 doublets and central pair discernible. Bottom-tilted view showing the region in which the axoneme bends and is no longer perpendicular to the imaging plane. This is associated with a decreased resolution not enabling to discern each doublet (arrowhead). E- Examples of axonemal regions of procyclic *T. brucei* cells analyzed for how much they deviated from circularity by determining the aspect ratio of the major and minor axis of a fitted ellipse. Sum of 5 consecutive z-planes is shown. For remaining analyzed regions see Suppl. fig. 5.

Dividing cells were readily identifiable by the presence of two axonemes. The distal end of the new axoneme was found in the proximity of the old axoneme with a gap separating the two (Fig. 1C, 1D, and Suppl. fig. 1). It had been shown that this space is occupied by the membrane junction termed the flagella connector (Moreira-Leite et al., 2001). Due to the ability to rapidly screen cells in a population at a low magnification, it was straightforward to identify and image rarer cell types, such as cells in various stages of cytokinesis (Figs. 1C, 1D, and Suppl. fig. 1B). In these cells the cleavage furrow was clearly discernible and the non-equivalence of nascent daughter cells apparent, with the new-flagellum daughter having a longer free flagellum overhang and a non-tapered anterior end (Fig. 1C, 1D, and Suppl. fig. 1B) (Abeywickrema et al., 2019).

Similarly to the procyclic cells, staining expanded bloodstream *T. brucei* cells with the anti-tubulin antibodies facilitated visualization of the shape of the corset and the axoneme (Fig. 1E). The increased resolution made discernible the area of missing anti-tubulin signal surrounding the distal end of the new axoneme (Fig. 1F). This corresponds to the groove, an indentation of the cell body membrane with an associated gap in the subpellicular corset originally observed by serial block-face scanning EM (Hughes et al., 2013); ExM enabled for the first time visualization of the groove by light microscopy. In addition to the groove, a line devoid of the signal extended alongside each axoneme; this area was likely occupied by the FAZ filament (Fig. 1F).

As *L. major* cells are of significantly smaller dimensions than *T. brucei* cells, their expansion prior to imaging would be very beneficial. Staining with the anti-tubulin antibodies indeed facilitated a 3D reconstruction of corsets and axonemes of expanded promastigote *L. major* cells. We observed slender cells with a single flagellum as well as shorter and rounder cells in cytokinesis (Fig. 1G and 1H), recapitulating the previously described changes to the cell shape associated with the cell cycle progression (Ambit et al., 2011; Wheeler et al., 2011). The expanded cells were between 30 - 60 µm long, which compared to non-expanded cells (6.5 - 13 µm) (Suppl. fig. 3) implicated the expansion factor of approximately 4.6, similar to the one observed for procyclic *T. brucei* cells.

**Figure 3.**
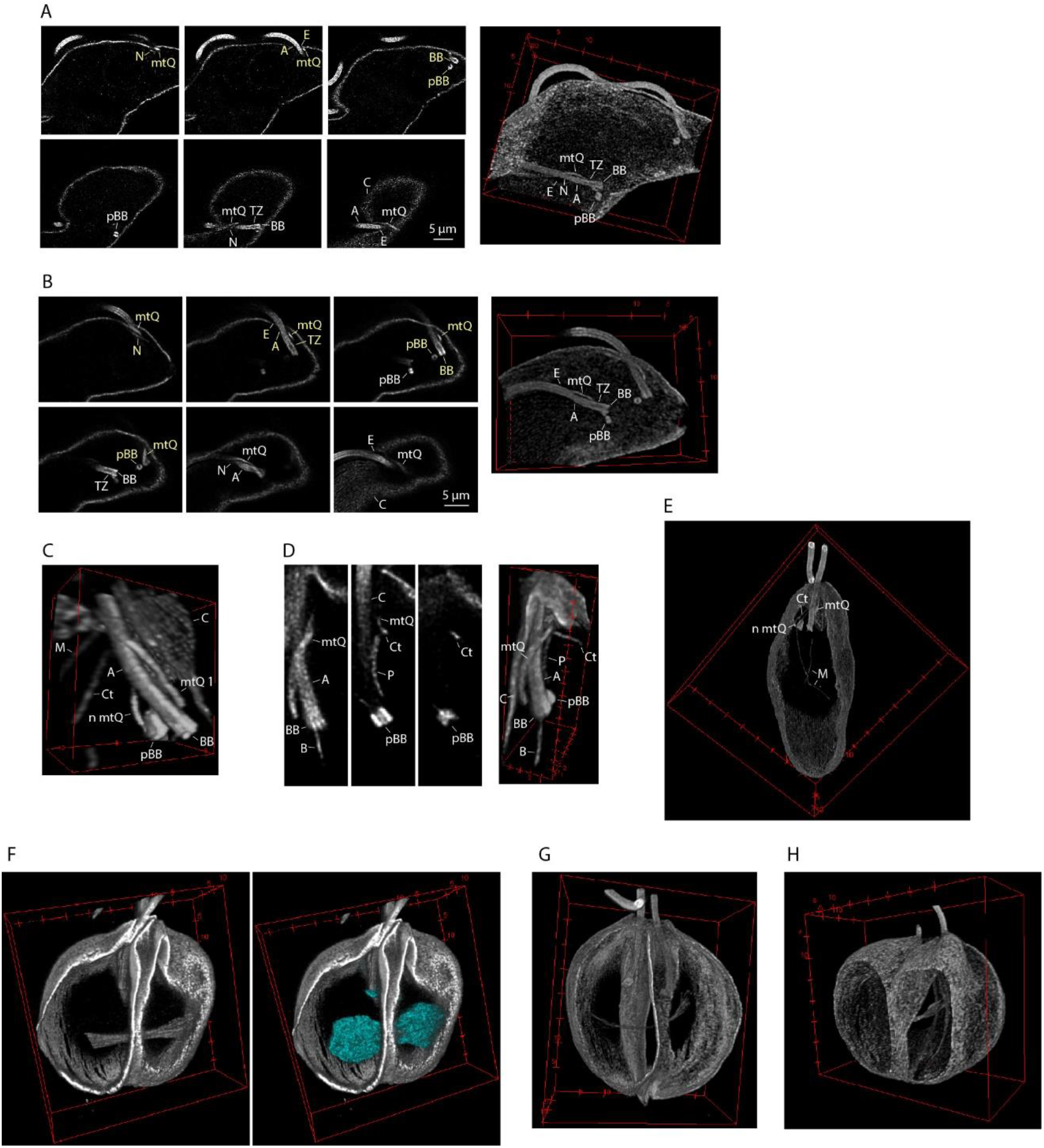
ExM facilitates studies of structures inside the corset. A- Individual z-planes (left) and 3D reconstruction (right) of the BB area of an expanded procyclic *T. brucei* cell (also shown in Fig. 1C) stained with C3B9. Yellow labels-structures associated with the new flagellum; white labels - structures associated with the old flagellum. A - axoneme; BB - basal body; pBB - pro-basal body; C - corset microtubules; mtQ - microtubule quartet; TZ - transition zone; E - exit of the flagellum from the corset; N - neck microtubule. Note that the mtQ signal is visibly broader than the signal of a single corset microtubule. B- Individual z-planes (left) and 3D reconstruction (right) of the BB area of an expanded bloodstream *T. brucei* cell (the cell shown in Fig. 1E) stained with C3B9. Yellow-structures associated with the new flagellum; white-structures associated with the old flagellum. A - axoneme; BB - basal body; pBB - pro-basal body; C - corset microtubules; mtQ - microtubule quartet; TZ - transition zone; E - exit of the flagellum from the corset; N - neck microtubule. Note that the mtQ signal is visibly broader than the signal of a single corset microtubule. C- 3D reconstruction of the BB area of an expanded *L. major* cell with a single flagellum (the cell shown also in Fig. 1G). The cell was stained with C3B9. A - axoneme; BB - basal body; pBB - pro-basal body; C - corset microtubules; mtQ - microtubule quartet; n mtQ - nascent mtQ associated with the pBB; Ct - cytoplasmic microtubules; M - part of a microtubules associated with the multivesicular tubule. D- Individual confocal z-planes (left) and 3D reconstruction (right) of the BB area associated with one of two flagella in an expanded mitotic *L. major* cell, which was stained with C3B9 (see also Fig. 3G). A - axoneme; BB - basal body; pBB - pro-basal body; C - corset microtubules; mtQ - microtubule quartet; P - pocket microtubules; Ct - cytoplasmic microtubules; B - individual microtubules extending from the BB or pBB. E- View inside the 3D reconstructed expanded *L. major* cell (identical to the cell in Fig. 1G) revealed long microtubules (M) likely corresponding to the previously described microtubules associated with the multivesicular tubule. Note also morphology of the cytoplasmic microtubule (Ct), and the nascent (n mtQ) and existing microtubule quartet (mtQ) in the BB area (shown in a higher magnification in Fig. 3C). F- H- View inside the 3D reconstructed corsets of expanded mitotic *L. major* cells. The mitotic spindle became thinner and elongated (from F to H), ultimately having flared microtubule ends at the spindle poles. The spindle was surrounded by DNA as revealed by Hoechst stain (cyan). Note also the cleavage furrow separating the cells into two daughters along their anterior-posterior axis and approximately perpendicular to the spindle.

3D reconstruction of cytoskeletons of cells in cytokinesis revealed that the cleavage furrow partitioned the cytoskeleton along the anterior-posterior axis of the cell body between the two flagella (Fig. 1H). The last point of attachment of two daughter cells was at their very posterior end (Fig. 1H), as was previously described by scanning EM (Wheeler et al., 2011) and at a lower magnification also by differential interference contrast microscopy (Ambit et al., 2011).

### 2/ ExM resolves individual structural components

Inspecting confocal planes revealed that in many individual corset microtubules could be discerned (Fig. 2A). This was particularly the case when the microtubules were oriented parallel to the imaging plane and close to the surface of the gel, with the concomitantly high signal intensity. To estimate the lateral spacing of the microtubules we performed the forward Fourier transform of images of 10 such regions (for an example see Fig. 2B). The resulting frequency domain images showed strongly represented frequencies with periodicities within an interval with boundaries of 270 ± 30 nm (mean ± SD) and 330 ± 50 nm, and with the mean value of 300 nm. These frequencies indeed accounted for the majority of observed microtubule-associated signal (Fig. 2B). The spacing of corset microtubules was previously determined by transmission EM as being 24 nm (Lacomble et al., 2009). Taking into account the microtubule diameter of about 25 nm, the centre to centre distance of two adjacent microtubules in an unexpanded cell is about 50 nm. This implicates an expansion factor of 6.0, significantly higher than the expansion factor of the gel (4.7).

The antibodies KMX-1, TAT1, and C3B9 produced staining of a higher density and lower background than the rabbit anti-tubulin antibody. However, using none of these anti-tubulin antibodies it was possible to reliably determine where individual microtubules terminated in corsets of *T. brucei* (e.g. Fig. 2A and 2B). This was, however, possible in *L. major*, mainly due to larger distances between corset microtubules in expanded cells (periodicities within an interval of 310 ± 20 nm to 470 ± 30 nm, n = 11 cytoskeletal regions; see also Suppl. fig. 4). This, for example, enabled visualization of the high concentration of microtubule ends at the posterior end of the cell (Fig. 2C).

**Figure 4.**
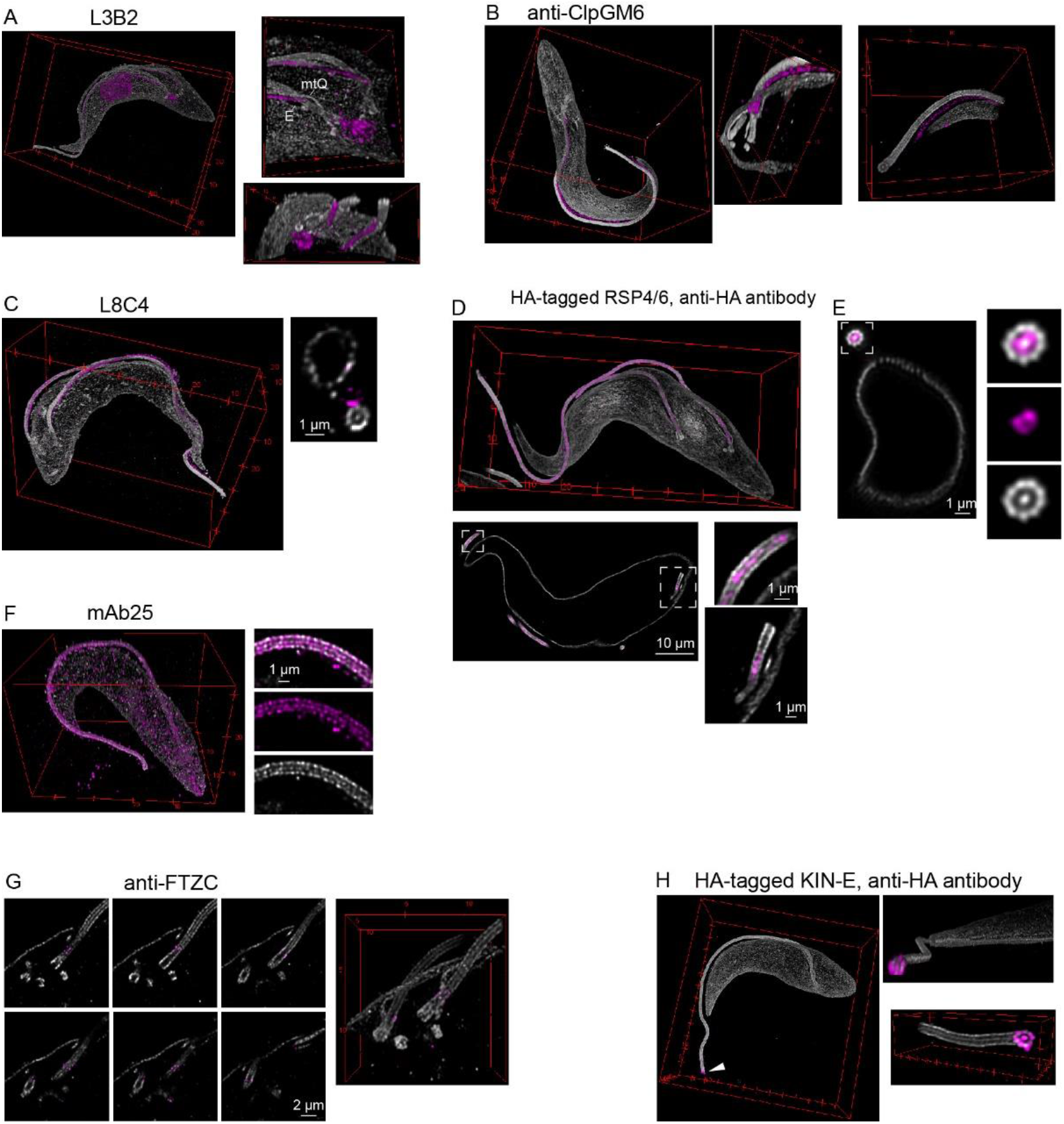
Localizing cytoskeletal proteins in respect to microtubule-based structures. Shown are expanded procyclic *T. brucei* cells, anti-tubulin signal shown in shades of gray, signal of antibodies to other cytoskeletal constituents in magenta. A- Confocal imaging of a cell stained with L3B2 and the polyclonal anti-alpha tubulin antibody. Left-3D reconstruction of the cellular volume; Right - en face (top) and from the anterior end (bottom) view of the BB area without the bottom most part of the microtubule corset. E - exit of the flagellum from the corset; mtQ - microtubule quartet. Note the nuclear and kinetoplast signals produced by the antibody. B- Confocal imaging of a cell stained with an anti-ClpGM6 antibody and C3B9 in a gel expanded in 84% sucrose. Left - 3D reconstruction of the cellular volume; Middle - a view of the 3D reconstructed BB area without the bottom and top most corset microtubules; Right - distal part of the axoneme with the anterior end of the corset of a cell different to the one shown on right, and imaged at a higher magnification. C- Confocal imaging of a cell stained with L8C4 and the polyclonal anti-alpha tubulin antibody. Left - 3D reconstruction of the cellular volume; Right - an orthogonal optical section through the anterior part of the cell. D- Confocal imaging of a cell expressing HA-tagged RSP4/6 and stained with the anti-HA antibody and with C3B9. Top - 3D reconstruction of the cell. Bottom - A single confocal plane of the cell. Insets show longitudinal optical sections through the axoneme and the proximal region of the flagellum. E- A single confocal plane of a cell (different to the cell in 4D) expressing HA-tagged RSP4/6 and stained with the anti-HA antibody and with C3B9. The inset shows an optical cross-section through the axoneme. The images were acquired using 8 x line averaging. F- Confocal imaging of a cell stained with mAb25 and the polyclonal anti-alpha tubulin antibody. Left-3D reconstruction of the cellular volume; Right - a single confocal plane of the axoneme - (top to bottom) merge, mAb25 signal, anti-tubulin signal. G- Confocal imaging of the BB area without the bottom and top most corset microtubules of a cells stained with FTZC and C3B9. Left - individual z-planes; Right - 3D reconstruction. H- Confocal imaging of a cell expressing 3 x HA-tagged KIN-E and stained with the anti-HA antibody and with C3B9. Left - 3D reconstruction of the cellular volume. Arrowhead indicates the distal end of the flagellum with the anti-HA antibody signal; Right top - the view from the distal end of the flagellum of the cell on left. Note that both images come from a single confocal imaging and were obtained at an identical magnification; Right bottom - Confocal imaging of the distal end of a flagellum of a cell different to the one shown on left and acquired at a higher magnification. Note the unambiguous localization of KIN-E to the ends of all doublets and the central pair. Tubulin signal of the same axoneme is shown also in Fig. 2D.

In individual confocal planes axonemal microtubule doublets were readily noticeable (Fig. 2A). In regions where the axoneme was oriented perpendicular to the imaging plane, it was possible to discern all 9 doublets and the central pair, which was typically seen as a unit (Fig. 2D). 3D reconstruction revealed that in other regions not all the doublets were resolved (Fig. 2D), owning to the significantly worse axial than lateral resolution in light microscopy (Tam and Merino, 2015).

Based on transmission EM data the axoneme is considered to be circular in its cross-section (Gadelha et al., 2006), enabling us to assess whether the expansion in ExM was isotropic. We analysed 27 axonemal regions of procyclic *T. brucei* specimens expanded either in water or in 84% sucrose, in which the axonemes were oriented perpendicular to the imaging plane. Fitting an ellipse to the axonemal cross-sections revealed that the aspect ratio between the major and minor axis was in all cases close to one (Fig. 2E and Suppl. fig. 5), i.e. the expanded axonemes were nearly circular. There was no significant difference between specimens expanded in water (1.099 ± 0.037, n = 9) and in sucrose (1.096 ± 0.044, n= 18).

Furthermore, diameter calculated from the circle area was 1.060 ± 0.022 µm^2^ for 9 specimens expanded in water, and 0.946 ± 0.010 µm^2^ for 18 specimens expanded in sucrose. When divided by the diameter of the axoneme (220 nm according to Pigino et al., 2011), it yielded the expansion factor of 4.8 in water and 4.3 in sucrose, well in accord with the measured expansion factors of gels (4.7 and 4.3, respectively).

### 3/ ExM facilitates studies of structures inside the corset

Intriguingly, ExM also facilitated visualization of tubulin-based structures located inside the corset. In both procylic (n = 38 analysed cells) and bloodstream (n = 14) *T. brucei* cells these were found in the basal body (BB) area around the proximal part of the flagellum. Their identity and organization became apparent when a 3D volume of the area was reconstructed (Fig. 3A and 3B), as there was a very good agreement with the previously published model based on electron microscope tomography (Lacomble et al., 2009).

The strongly labelled BB could be distinguished at the base of the flagellum. The BB transited via a hollow transition zone into the axoneme, which was identifiable by the presence of the central pair. The axoneme exited the corset via a space visibly devoid of the microtubules. The pro-basal body (pBB) was located adjacent to each BB, and its orientation could be determined. Furthermore, a ribbon-like structure was spiralling alongside the proximal part of each axoneme. The structure started on the side of the BB facing the pBB and extended towards the corset, with which it merged. The position, shape, and the fact, that it was stained by various anti-tubulin antibodies (Suppl. fig. 6) were consistent with the structure being the microtubule quartet (mtQ), which subtends the flagellar pocket (Lacomble et al., 2009). The mtQ is typically formed by 4 microtubules (Lacomble et al., 2009; Sherwin and Gull, 1989). Although in ExM images the mtQ signal was substantially wider than the signal of a corset microtubule (Fig. 3A and 3B), it was not possible to resolve individual mtQ microtubules. This is consistent with the mtQ microtubules in the basal body area being more tightly spaced than the corset microtubules (e.g. Fig. 3B in Höög et al., 2012).

Interestingly, in expanded cells a single microtubule extending alongside the mtQ but terminating before the mtQ integrated into the corset was noticeable (Fig. 3A, 3B, and Suppl. fig. 6), which corresponded to the neck microtubule previously described by electron microscope tomography (Lacomble et al., 2009).

In expanded procyclic cells this neck microtubule was observed in 54 of 56 BB areas (associated with the single flagellum of non-dividing cells, the old flagellum in dividing cells with two flagella, or the new flagellum in dividing cells, which extended significantly past the exit point). Similarly, the neck microtubule was present in 21 of 22 BB areas in bloodstream cells (for an example, see Fig. 3B).

EM previously described an even higher complexity of the microtubule-based structures inside the corset of *Leishmania* (Weise et al., 2000; Wheeler et al., 2016). Our ExM data were well in accord with this. Moreover, the quantitative aspect of ExM revealed a significant variability in the presence of some of these structures in a population of promastigote *L. major* cells.

We analysed 3D reconstructed volumes of 7 cells with a single flagellum and 17 cells with two flagella. All 41 axonemes present in these cells were subtended by the BB with the associated pBB (Fig. 3C and 3D). Moreover, all the axonemes had the mtQ extending alongside their proximal part (Fig. 3C and 3D). Unlike in *T. brucei*, the mtQs did not intercalate the corset (Fig. 3C and 3D), confirming the EM observations (Wheeler et al., 2016). Although the signal associated with the structure in ExM was wider than a single corset microtubule (Fig. 3C and 3D), individual mtQ microtubules could not be resolved. This is consistent with smaller distances between microtubules in the mtQ than in the corset (see Fig. 4E in Weise et al., 2000; and Fig. 3D in Wheeler et al., 2016). In 4 of the 7 cells with a single flagellum a microtubule-based structure extending in an arch from the pBB towards the axoneme was present (Fig. 3C and 3E); it likely corresponded to the nascent mtQ previously observed by EM in this cell type (Wheeler et al., 2016).

Additional structures included pocket microtubules, observed alongside 20 axonemes on the opposite side to the mtQ (Fig. 3D), and cytoplasmic microtubules, which were also associated with 20 axonemes (only partially overlapping with the set of axonemes with the pocket microtubules), but ran orthogonal to the mtQ (Fig. 3C and 3D). Finally, we observed short microtubules or small plate-like structures attached to 14 of the 41 BBs or associated pBBs (Fig. 3D). It is worth noting that in a minority of cases morphology or positioning of these additional structures deviated from those described by EM, highlighting the significant degree of variability of the microtubule-based structures in the BB area of *Leishmania*. This, together with the lack of other positional information, such as the flagellar pocket membrane, in the ExM specimens, made in these cases classification of the structures challenging.

Intriguingly, inside the corset of 20 of the 24 reconstructed cells, one (n = 17) or two (n = 3) long microtubules were present. These microtubules were less straight than microtubules of the corset and extended from its anterior end to the central part (Fig. 3C and 3E). Occasionally, an entire microtubule was located in the central part (Fig. 3E). These microtubules corresponded to the microtubules associated with the multivesicular tubule observed by EM in cell sections (Weise et al., 2000). Interestingly, all 4 cells, in which no multivesicular tubule-associated microtubules were present, were in mitosis. However, in other mitotic cells these microtubules were observed.

In mitotic *Leishmania* cells ExM visualized well the mitotic spindle, which was formed by a bundle of microtubules (Fig. 3F), as had been previously described by transmission EM of serial sections (Ureña, 1986). In expanded cells individual spindle microtubules were discernible, particularly at spindle poles. DNA staining demonstrated that the spindle was surrounded by the nuclear DNA, segregated by the spindle into two masses (Fig. 3F). Cells at later stages of mitosis possessed thin elongated spindles with flared microtubule ends localized at the poles (Fig. 3G and 3H). In observed mitotic cells an ingression of the corset microtubules along the anterior-posterior cell axis was present (Fig. 3F - 3H). The cleavage plane was orthogonal to the axis connecting the BBs of the two flagella and to the long axis of the mitotic spindle (Fig. 3G and 3H).

### 4/ Localizing cytoskeletal proteins in respect to microtubule-based structures

ExM has been frequently used to determine position of various proteins in respect to microtubule-based structures (e.g. Amodeo et al., 2021; Bertiaux et al., 2021; Halpern et al., 2017; Le Guennec et al., 2020).

Procyclic *T. brucei* cells, due to their well-defined morphology, a considerable body of knowledge on protein localization acquired by EM, and availability of high-quality antibodies (see Suppl. fig. 2) appear to be an excellent system to further validate the approach. First, we focused on FAZ and assessed whether ExM can differentiate proteins localizing to distinct domains of the structure. Staining expanded cells with the L3B2 antibody recognizing the FAZ1 protein (Kohl et al., 1999; Vaughan et al., 2008) produced a linear signal starting below the point of the axoneme exiting the corset and intimately linked to the mtQ (Fig. 4A). It further extended along the corset following the axoneme and terminated at the anterior end of the corset (Fig. 4A). Importantly, the signal was positioned between microtubules of the corset and also extended below its surface (Fig. 4A). This is consistent with the localization of the FAZ1 protein to the FAZ filament domain in the cell body, as determined by immunogold EM (Kohl et al., 1999; Sunter and Gull, 2016). In contrast to immunofluorescence microscopy, the antibody also stained the nucleus and the kinetoplast in ExM (Fig. 4A and Suppl. fig. 2).

Another FAZ protein, ClpGM6, was previously shown by immunogold EM to constitute FAZ fibres in the FAZ flagellum domain linking the flagellar cytoskeleton to the membrane junction (Müller et al., 1992). Staining expanded cells with the anti-ClpGM6 antibody (Hayes et al., 2014) indeed produced a robust signal spanning the gap between the axoneme and the corset (Fig. 4B). The signal started around the point of the axoneme exiting the corset and terminated past its anterior end (Fig. 4B).

Next, we focused on individual structures of the flagellar cytoskeleton. First, we applied to expanded cells the L8C4 antibody, which recognizes a constituent of the paraflagellar rod (Kohl et al., 1999), a kinetoplastid-specific structure extending alongside the axoneme. In ExM the L8C4 signal was indeed juxtaposed to the axoneme (Fig. 4C). Despite its significant granularity it was possible to determine that the signal started distal to the point of the axoneme exiting the corset and diminished towards the distal end of the axoneme, which is consistent with immunofluorescence observations (Suppl. fig. 2; see also Kohl et al., 1999).

As an example of a protein found inside the axoneme we studied RSP4/6, an evolutionary conserved member of the radial spokes connecting the microtubule doublets to the central pair (Pigino et al., 2011). As no antibodies against the trypanosome RSP4/6 are available, we tagged the protein at its endogenous locus with a 3 x HA tag and stained the resulting cells after expansion with an anti-HA antibody. In both, optical cross-sections and longitudinal sections it was apparent that the anti-HA signal was confined within the space defined by the outer doublets (Fig. 4D and 4E). Moreover, in the orthogonal sections it was noticeable that the central part was devoid of the signal; this corresponded to the space occupied by the central pair microtubules (Fig. 4E). ExM also revealed that the signal started just distal to the TZ concomitant with the central pair microtubules (Fig. 4D).

Next, we studied localization of the protein *Tb*SAXO. This microtubule-associated protein had previously been demonstrated to localize to the axoneme by immunogold EM (Dacheux et al., 2012) and has been used as an axonemal marker in *T. brucei*. Expanded cells stained with the mAb25 antibody (Pradel et al., 2006) recognizing *Tb*SAXO revealed a clear co-localization of the protein with tubulin in the axoneme (Fig. 4F). A further interpretation of the mAb25 staining, such as the point at which the signal started along the axoneme, was precluded by the relatively strong speckled background staining in and around the cell.

Finally, we assessed the level of positional information achievable by ExM for proteins localizing to discrete regions along the flagellar cytoskeleton. First, we stained expanded cells with an antibody against FTZC, the first described marker of the transition zone in trypanosomes (Bringaud et al., 2000). ExM revealed that individual puncta of the signal surrounded microtubules of the transition zone (Fig. 4G). Next, we generated a procyclic cell line expressing 3 x HA-tagged KIN-E, the protein localizing along the flagellum and significantly enriched at the flagellum distal end (An and Li, 2018). In expanded cells an anti-HA antibody indeed produced a robust staining at the very end of the axoneme (Fig. 4H). Intriguingly, the resolution provided by ExM allowed to unambiguously localize KIN-E to the distal end of each axonemal doublet as well as to the central pair (Fig. 4H).

## Discussion

In this work we set out to assess the ExM approach for studying cytoskeletal structures using unicellular kinetoplastid parasites. Morphology of their cytoskeleton, particularly in the case of procyclic and bloodstream *T. brucei* cells, had been extensively characterized by various EM approaches, with a remarkably little variation between cells in a population. We observed that ExM is capable of faithfully reproducing information on the presence, positioning, and morphology of structures and structural components. A prime example being the mtQ, which in 3D reconstructed cellular volumes obtained by ExM had an identical position, helical pattern and handedness as those previously determined by EM tomography (Lacomble et al., 2009). Other notable examples include the groove of the bloodstream cells, so far observed only by serial block-face scanning EM (Hughes et al., 2013), and differential localization of proteins to individual domains of the FAZ, previously determined by immunogold EM (Kohl et al., 1999; Müller et al., 1992). Hence, ExM can serve as a substitute of these EM approaches, which require highly specialized instrumentation, when the lateral resolution of about 50 nm suffices.

Our analysis of the axoneme shape showed that the expansion was isotropic, which is consistent with published observations of tissues, cells, organelles, and structures (Chen et al., 2016; Gambarotto et al., 2019; Tillberg et al., 2016). Our data, however, also indicated that the degree of expansion was not identical for the studied cytoskeletal structures; while the axoneme expanded 4.8 fold, similar to the 4.7 expansion factor of the gel, the distance between the corset microtubules increased 6.0 fold. Leaving aside the potential caveats associated with estimating dimensions from EM data (Shah et al., 2015), which were used to calculate these expansion factors, similar and higher discrepancies in expansion factors of individual organelles were recently observed in various experimental systems and using a variety of ExM protocols (Büttner et al., 2020; Kubalová et al., 2020; Pernal et al., 2020; Pesce et al., 2019). Hence, it seems likely that based on their composition individual organelles and structures undergo differential expansion, precluding a straightforward interpretation of measured distances between entities of expanded specimens.

Despite these limitations there are characteristics which make ExM well suited to complement EM approaches for answering specific questions.

i/ Foremost, ExM is based on immunofluorescence staining and standard confocal microscopy, with image acquisition and subsequent 3D reconstruction being comparatively rapid. Hence, it is straightforward to obtain 3D volumes of tens of cells, as presented here, with even hundreds of cells manageable. This would be challenging with EM approaches and facilitates quantitative super-resolution studies of cytoskeletal structures. Examples presented in this study include the observation of the neck microtubule being an integral part of the BB area in procyclic and bloodstream *T. brucei* cells, and that the mtQ is found alongside every axoneme in promastigote *L. major* cells, while pocket microtubules, cytosolic microtubules, and multivesicular tubule-associated microtubules are common, but not universally present.

ii/ The real strength of ExM lies in the method offering the possibility of rapidly inspecting a population of cells and at the same time the ability to image an entire volume of a cell of interest at a relatively high resolution. This unique combination makes possible studies of rare cell types, such as cells in transient phases of the cell cycle, for example in cytokinesis (Fig. 1C, 1D, and 1H). In these cells ExM visualized how the corset was partitioned by the cleavage furrow. In *L. major* ExM also provided information on morphology of the mitotic spindle, including observations of ends of individual microtubules at the spindle poles (Fig. 3F - 3H). In addition, ExM has an obvious potential to provide novel information on events leading to duplication of structures in the BB area and their re-organizations following initiation of the new flagellum construction (for example see Fig. 3C and 4B). Finally, we envisage that ExM will be in a similar way highly useful to study morphology and cellular architecture of various transient life cycle forms and the processes of cell differentiation.

iii/ Protein localization by immunogold EM is challenging as antibodies frequently do not reach or recognize their epitopes in resin-embedded sections. In contrast, it has been observed that antibody markers established for the standard immunofluorescence staining generally also stain expanded specimens (Ku et al., 2016). Our data are in line with this (see Table 1). Intriguingly, we observed that in expanded cells the anti-tubulin antibodies stained the axonemal microtubules well, while they have been known to produce only a weak signal in non-expanded cells (Fig. 1A - 1C and Suppl. fig. 2; see also Woods et al., 1989). This likely reflects an increased accessibility of their epitopes in the axoneme due its expansion. Such a positive effect of de-crowding due to ExM was previously described in other systems (Sarkar et al., 2020; Tillberg et al., 2016). On the other hand, we observed that two tested antibodies, BBA4 staining the BBs and pBBs in *T. brucei* (Woods et al., 1989) and mAb62 staining the flagella connector (Abeywickrema et al., 2019) (see also Suppl. fig. 2), did not produce any specific signal in ExM (Suppl. fig. 7). This may stem from a degradation of respective epitopes during the SDS-based denaturation step of ExM, not present in a standard immunofluorescence protocol. An ExM approach based on a different homogenization step could be considered (Chozinski et al., 2016).

The applicability of ExM to localize proteins in these kinetoplastid species is further augmented by the available repertoire of reverse genetics approaches. We used the rapid PCR-only approach to generate cell lines expressing proteins of interest tagged at their endogenous loci with a small epitope tag (Dean et al., 2015). The resolution of ExM images was sufficient to unambiguously localize the radial spoke protein RSP4/6 to the space between the axonemal microtubule doublets and the central pair, and the kinesin KIN-E to the tips of individual doublets as well as the central pair. Hence, this approach opens up the possibility of localizing in the cellular volume and at a relatively high resolution many proteins encoded in genomes of these parasites.

In conclusion, ExM is a powerful approach for studying cytoskeletal structures in *T. brucei, L. major*, and likely other kinetoplastid species. It is a simple, yet robust super-resolution method, which does not require highly specialized reagents or instrumentation. ExM data recapitulate well observations by various EM approaches. Moreover, ExM is capable of complementing them due to its ability to provide a highly quantitative information.

## Acknowledgment

We are grateful to Farnaz Zahedifard and Martin Zoltner for providing bloodstream *Trypanosoma brucei brucei* cells and promastigote *Leishmania major* cells. We thank Marketa Schmidt-Cernohorska for introducing us to expansion microscopy, Ludek Stepanek for assistance with image analysis, and Keith Gull for providing antibodies and for critical comments on the manuscript. Antibodies were also provided by Derrick Robinson and Frederic Bringaud. Work in the Laboratory of cell motility was supported by the Czech Science Foundation (GA CR) project no. 20-23165J and by an Installation Grant from the European Molecular Biology Organization. V.V. is a holder of the J. E. Purkyne Fellowship of the Czech Academy of Sciences. The study was also supported by the Charles University project GA UK No. 972120 to P.G. We further acknowledge the Light Microscopy Core Facility, IMG CAS, Prague, Czech Republic, supported by MEYS (LM2018129, CZ.02.1.01/0.0/0.0/18_046/0016045) and RVO: 68378050-KAV-NPUI, for their assistance with the confocal imaging presented in this study, and Imaging Methods Core Facility at BIOCEV, institution supported by the MEYS CR (Large RI Project LM2018129 Czech-BioImaging) and ERDF (project No. CZ.02.1.01/0.0/0.0/18_046/0016045) for their support with data processing.

## Materials and Methods

### Cell Growth and Preparation of Transgenic Cell Lines

*Trypanosoma brucei brucei* procyclic SmOxP927 cells (Poon et al., 2012) were cultured at 28°C in SDM-79 medium (Life Technologies) supplemented with 10% fetal bovine serum (Gibco, 10270106) (Brun and Schönenberger, 1979). Transgenic cell lines with flagellar proteins KIN-E (Tb927.5.2410; An and Li, 2018) and RSP4/6 (Tb927.11.4480; Aslett et al., 2010) tagged with a 3 x HA tag were generated using the pPOT tagging approach (Dean et al., 2015). Bloodstream *Trypanosoma brucei brucei* cells of the 427 cell line and promastigote *Leishmania major* cells were provided by the laboratory of Martin Zoltner, Charles University, Prague, Czech Republic.

### Expansion Microscopy

#### Fixation of cells

For fixation on coverslips 2 x 10^6^ cells per specimen were used, and for fixation in solution 1 x 10^6^ cells per specimen were used. The cells were pelleted by centrifugation for 5 min at 800 g, followed by two washes with PBS, or with PBS supplemented with 3.3 mM glucose (Penta, G01102) in the case of bloodstream *T. brucei* cells.

For fixation on coverslips the cells were re-suspended in 50 µl of PBS and transferred onto a coverslip (High Precision ø 12 mm No. 1.5H, Marienfeld) placed in a 24-well plate (Jet Biofil, TCP011024), to facilitate subsequent steps. Following a 5 min incubation the cells were washed with PBS and fixed with 500 µl of a freshly prepared fixation solution containing 4% formaldehyde (Sigma, F8775) and 4% acrylamide (Sigma, A8887) in PBS. For fixation in solution the cells were directly re-suspended in the fixation solution and transferred to a poly-L-lysine (Sigma, P4707) coated coverslip in a 24-well plate. In both cases the cells were fixed overnight at room temperature, followed by a PBS wash.

#### Gelation

Gelation was performed on a strip of Parafilm in a wet chamber, which was initially placed on ice. The monomer solution containing 19% sodium acrylate (Sigma, 408220), 10% acrylamide, and 0.1% N, N’-Methylenebisacrylamide (Sigma, M7256) in PBS was aliquoted and stored at −20°C for up to a month. Immediately prior to gelation N, N, N’
s, N’-tetramethylethylenediamine (Sigma, T9281) and ammonium persulfate (Thermo Scientific, 17874) were added to the monomer solution to the final concentration of 0.5% each, and 50 µl of the mixture were quickly transferred onto Parafilm in the wet chamber. Coverslips with the cells facing down were placed on the drop of the monomer solution and incubated for 5 min. The wet chamber was subsequently transferred to 37°C and incubated for 30 min to allow for a proper gel polymerization.

#### Denaturation and expansion of the gel

The gelation chamber was brought to room temperature and several drops of ultrapure water were added to the coverslips to help the gel detach from Parafilm. Thereafter, the specimens were transferred to a well of a 12-well plate (Jet Biofil, TCP011012) containing 1 ml of the denaturation buffer consisting of 50 mM tris(hydroxymethyl)aminomethane-hydrochloride (Serva, 3719202), 200 mM sodium chloride (Lach-Ner, 30093-AP0), 200 mM sodium dodecyl sulphate (Roth, 1057), pH 9.0. At this point, gels started to expand, which facilitated their separation from the coverslips. Gels together with the remaining denaturation buffer were transferred to a 1.5 ml microcentrifuge tube, which contained 0.5 ml of the denaturation buffer, and denaturation was performed for 1 to 2 hours at 95°C. The denatured gels were transferred to ø 10 cm Petri dishes, and expanded by three 20 min incubations with 15 ml of ultrapure water.

#### Antibody staining of expanded specimens

Water was removed and an approximately 10 x 10 mm piece of the gel was cut per each antibody staining. Staining was performed in a 6-well plate (Jet Biofil, TCP011006), in the dark while gently rocking. Both primary and secondary antibodies were diluted in 500 µl of the ExM blocking buffer consisting of 2% bovine serum albumin (Roth, 8076) in PBS. In general, we observed that both primary and secondary antibodies produced good results when used more concentrated compared to standard immunofluorescence experiments. For antibody concentrations see Table 1. The gels were first incubated overnight at room temperature with 500 µl of diluted primary antibodies. Thereafter, they were washed by 3 x 20 min incubations with 3.5 ml of ultrapure water, followed by an overnight incubation with 500 µl of secondary antibodies. Subsequently, the gels were washed 3 x 20 min with 3.5 ml of ultrapure water and selected specimens were also incubated for 30 min at room temperature with 10 µg/ml 4′,6-diamidino-2-phenylindole (DAPI) (Sigma, D9542) or 10 µg/ml Hoechst 33342 (Thermo Scientific, 62249) in PBS while rocking. Finally, the gels were washed 5 x 15 min with 4 ml of ultrapure water. For co-staining experiments specimens were simultaneously incubated with two primary or two secondary antibodies.

#### Imaging

Water was carefully aspirated, a stained piece of the gel was transferred to a centre of a glass-bottom dish (Cellvis, D35-20-1.5-N) coated with poly-L-lysine, and a small drop of ultrapure water was put on it to prevent drying. To facilitate imaging it was important to keep track of the orientation of the gel during the procedure, such that the side of the gel with cells close to its surface was ultimately facing the glass-bottom dish. Specimens were imaged using a Leica TCS SP8 confocal microscope with an HC PL apochromatic 63x/NA 1.40 oil immersion objective. Excitation was performed with a 405 nm diode laser (50mW) in the case of DAPI and Hoechst 33342; a 488 nm solid-state laser (20 mW) in the case of Alexa Fluor 488, and a 552 nm solid-state laser (20 mW) in the case of Alexa Fluor 555. Fluorescence was detected using hybrid detectors (HyD). Z-stacks were acquired with the step size of 100 nm. The pixel size was between 52 - 97 nm, and in the case of high magnification images between 18 - 36 nm. The pixel dwell time was 400 ns. The size of the pinhole was adjusted based on the signal strength, but was typically around 0.4 Airy unit to improve resolution. No averaging was used for z-stacks. In the case of a significant lateral drift, a piece of a gel was embedded in the two-component silicone-glue Twinsil (Picodent, 13001000). Expanded gels could be stored in Twinsil for several days before imaging.

### Standard immunofluorescence staining

Procyclic *T. brucei* and promastigote *L. major* cells were washed and re-suspended in PBS to 1 x 10^7^ cells/ml. 50 µl of *T. brucei* cells were transferred onto a cleaned ø 12 mm coverslip, followed by a 4 min incubation. After removal of an excessive liquid the cells were fixed for 5 min with 4% formaldehyde in PBS, washed 3 x 5 min with PBS, and permeabilized for 10 min with 50 µl of 0.1% Triton X-100 (Roth, R30512) in PBS. 50 µl of *L. major* cells were transferred onto a cleaned microscopic slides (Menzel Gläser, 630-1985), incubated for 4 min, and an excessive liquid removed. The slides were transferred to an ice-cold methanol, incubated for 20 min at −20°C, and rehydrated for 30 min in PBS. Both *T. brucei* and *L. major* specimens were subsequently washed for 3 x 5 min with PBS and incubated for 1 hour with the IF blocking buffer consisting of 1% bovine serum albumin in PBS. Following another 3 x 5 min wash with PBS they were incubated for 1 hour with primary antibodies diluted in the IF blocking buffer. Then, the specimens were washed for 3 x 5 min with PBS, incubated for 1 hour with secondary antibodies diluted in the IF blocking buffer, washed, mounted into a mounting medium containing 90% glycerol with 1,4-diazabicyclo[2.2.2]octane and 100 ng/ml of DAPI, and stored at 4°C overnight. The specimens were subsequently imaged using a Zeiss Axioplan 2 microscope with a Zeiss 100x (NA 1.4) Plan Apochromat oil immersion objective and an Andor Zyla 4.2 camera controlled by Micromanager software (Edelstein et al., 2010).

### Image analysis

Confocal z-stacks and individual images of non-expanded cells were processed and analyzed in Fiji (Schindelin et al., 2012). 3D reconstructions were performed in the 3D viewer plugin. To calculate the Fourier transform of a 2D projected z-stack and to display the power spectrum the FFT function was used. To obtain images containing or omitting selected frequencies the Inverse FFT function was used. Selected z-stacks were deconvolved with Huygens Professional version 19.10 (Scientific Volume Imaging, The Netherlands, http://svi.nl), using the CMLE algorithm.

#### Assessing isotropy of expansion and extent of axonemal expansion

Isotropy of expansion was assessed on raw images (without deconvolution) based on the circular shape of the axonemal cross-section. First, axonemal regions perpendicular to the image plane were selected to avoid regions of elongated cross-sections due to the axoneme being tilted. The perpendicularity check was done in Fiji (Schindelin et al., 2012). The default threshold method was applied to manually selected series of z-planes with a single axoneme present to create an axoneme binary mask. The close operation was performed on the binary mask and the analyze particle tool was used to find the position of the mask centroid. For every five consecutive planes, the positions of the first and the fifth centroid were compared; if their xy coordinates were close to each other, they were further analyzed. The threshold for the xy centroid distance was chosen as 4 x z-step size/tan(85 degrees), which corresponded to a 5 degree tilt. 27 selected axonemal stacks of five z-planes were corrected for the drift (StackReg plugin based on Thevenaz et al., 1998, Rigid Body transformation), summed (sum slices), and the default threshold method was used to create a binary mask. Circularity of the mask was assessed using the analyze particle tool by fitting an ellipse to the mask and extracting the aspect ratio of its major to minor axis. The aspect ratio of one indicates a circular object.

The analyze particle tool provided also the area of the ellipse. A diameter of a circle with equivalent area was calculated. The diameter was corrected for blur due to light diffraction, estimated to be 200 nm (the value was obtained by distilling a point spread function in Huygens software from images of 100 nm beads acquired under conditions identical to z-stacks of cells). The corrected diameter divided by the typical diameter of the axoneme (220 nm according to Pigino et al., 2011) yielded the expansion factor.

---

**Supplementary figure 1.**
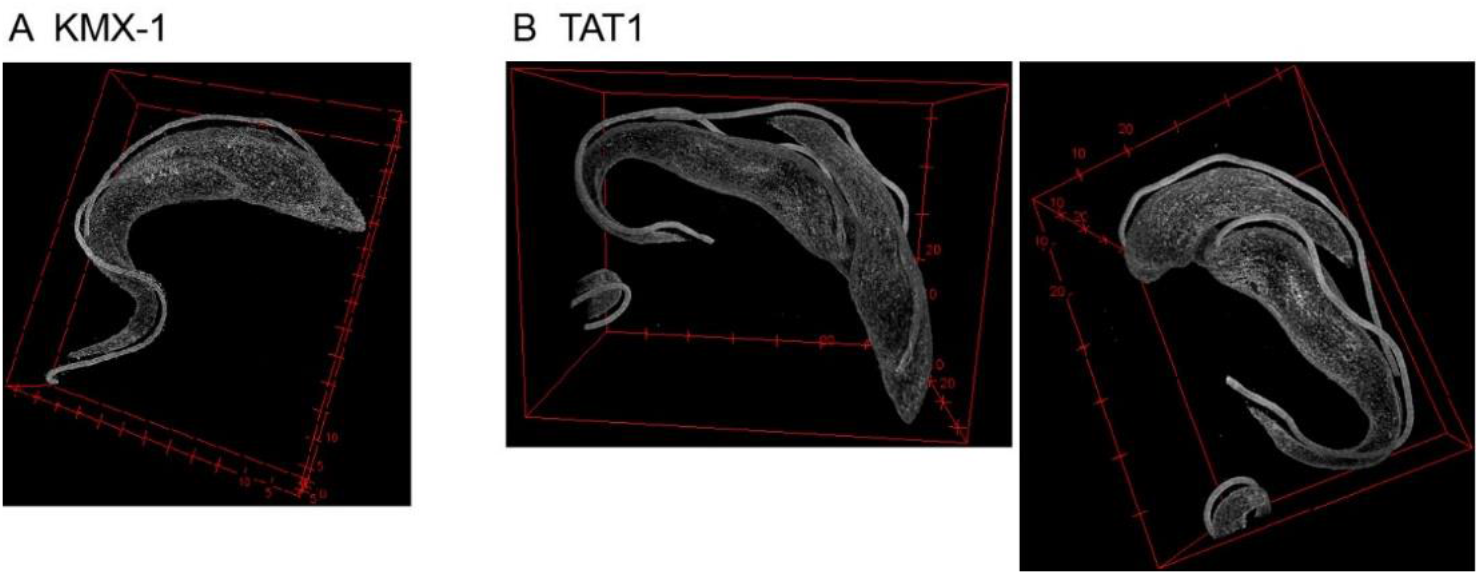
A - 3D reconstruction of a confocal z-stack of an expanded procyclic *T. brucei* cell fixed in solution before being settled on a glass coverslip and stained with the KMX-1 antibody. B - 3D reconstruction of a confocal z-stack of an expanded procyclic *T. brucei* cell fixed in solution before being settled on a glass coverslip and stained with the TAT1 antibody. Views from two different angels are shown to demonstrate morphology of the clevage furrow and non-equivalence in old- and new-flagellum daughter cells.

**Supplementary figure 2.**
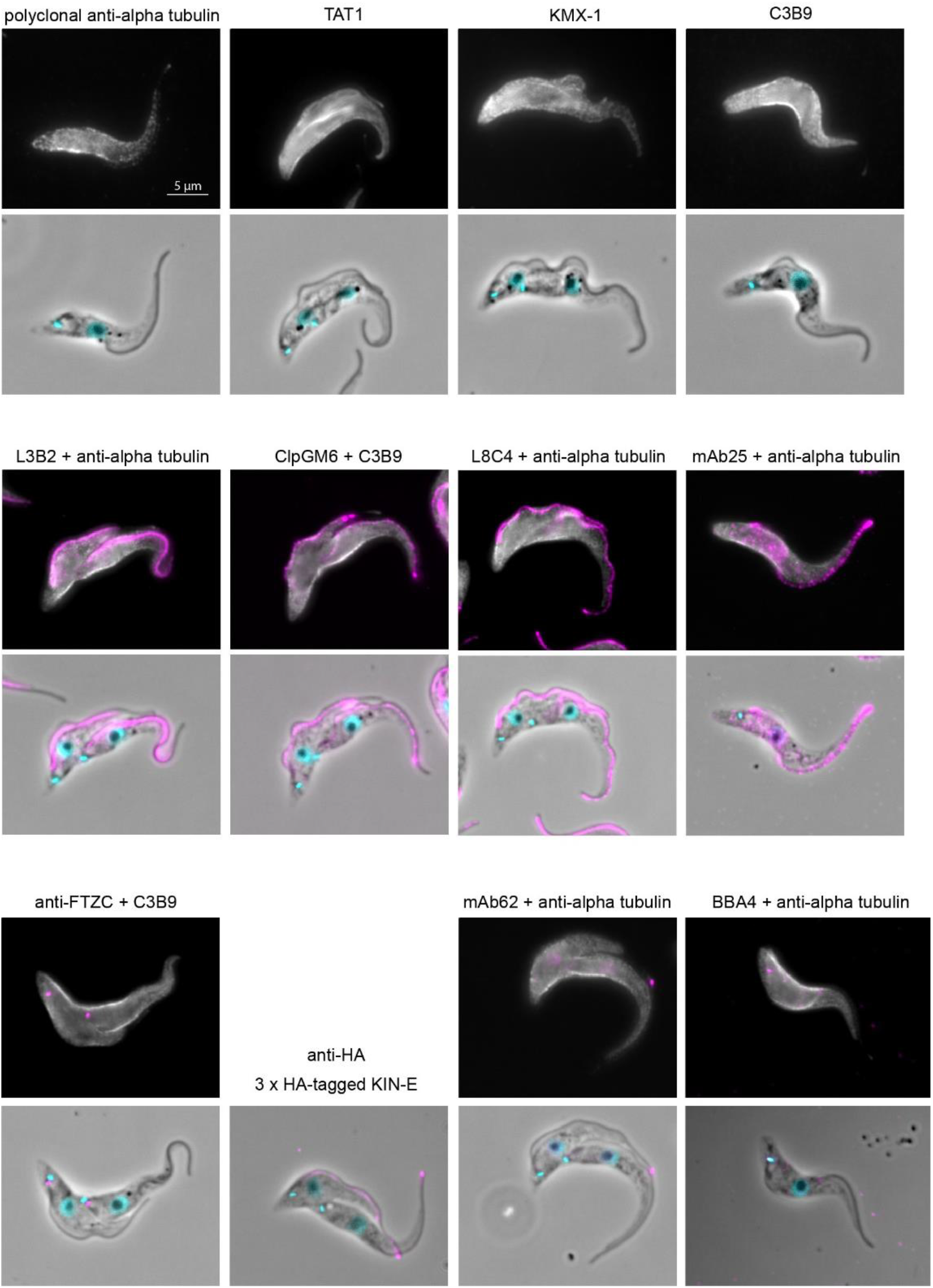
Immunofluorescence staining of non-expanded procyclic *T. brucei* cells. Signals of anti-tubulin antibodies (the polyclonal anti-alpha tubulin antibody, KMX-1, TAT1 or C3B9, top images) are in grey, signals of other used antibodies in magenta. Bottom images also show phase contrast (grey) and signal of the DNA stain DAPI (cyan).

**Supplementary figure 3.**
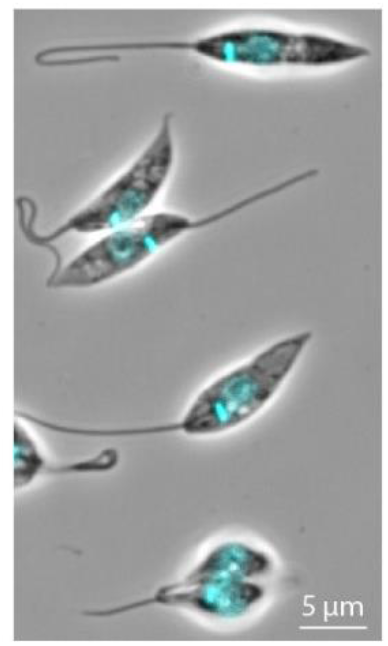
A phase contrast image of non-expanded promastigote *Leishmania major* cells merged with the signal of the DNA stain DAPI (cyan).

**Supplementary figure 4.**
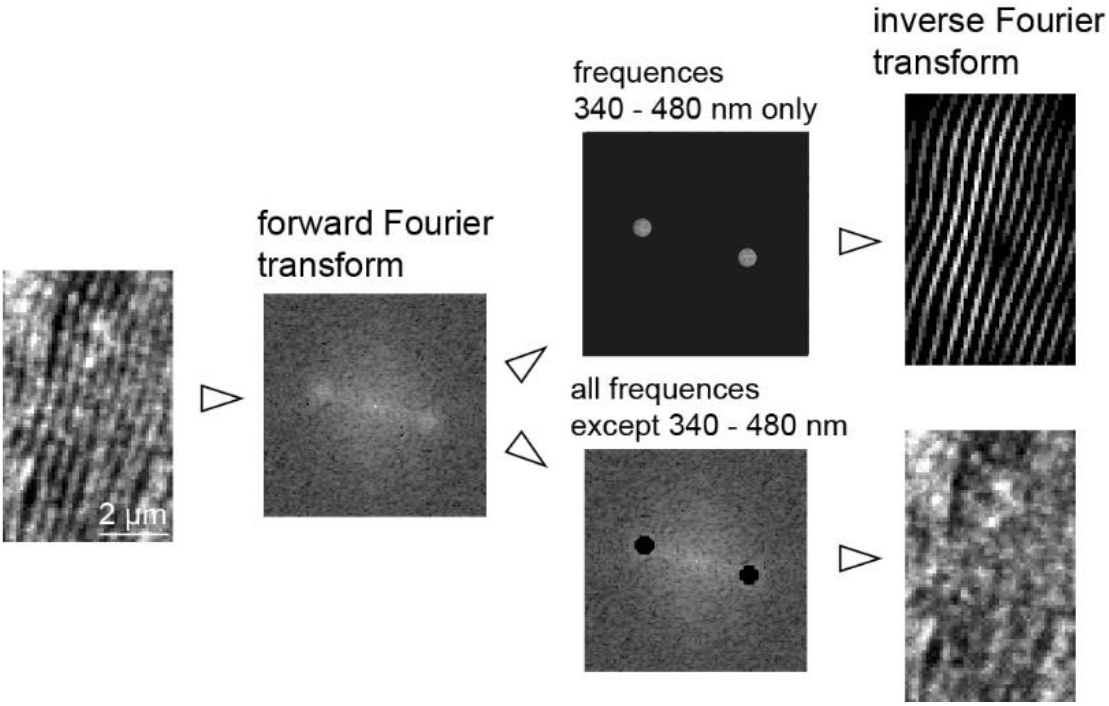
An example of an analysis of distances between microtubules in the subpellicular corset of *L. major*. Confocal z-planes encompassing corset microtubules in the vicinity of the glass surface and oriented approximately parallel to the image plane were projected to 2D by the average intensity algorithm. The forward Fourier transform of the resulting image was performed. The frequency domain image shows strongly represented frequencies with the periodicity of 340 - 480 nm. The inverse Fourier transform of these frequencies recapitulated the majority of the microtubule-associated signal (top), while the inverse Fourier transform of remaining frequencies did not (bottom).

**Supplementary figure 5.**
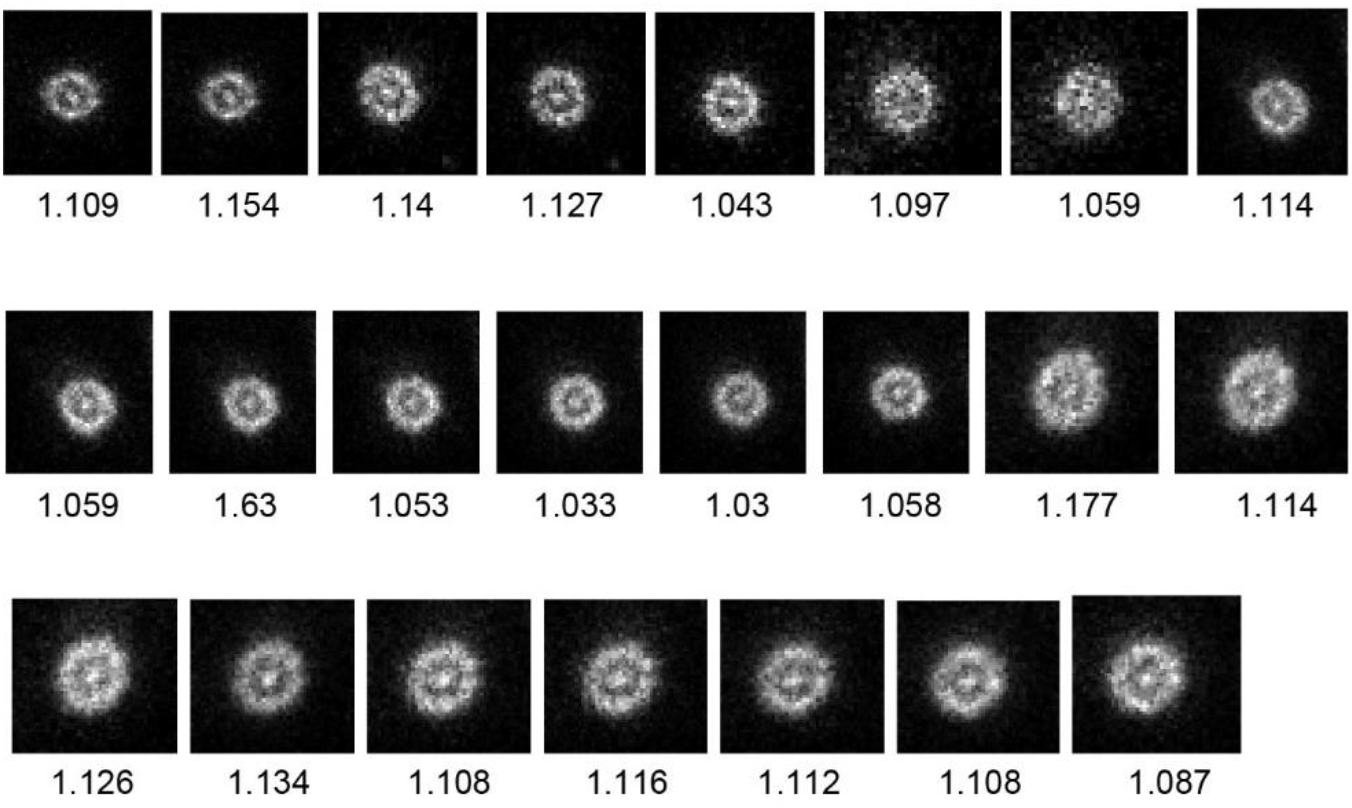
Axonemal regions analyzed for how much they deviated from circularity by determining the aspect ratio of the major and minor axis of a fitted ellipse (underneath each image). Sum of 5 consecutive z-planes is shown.

**Supplementary figure 6.**
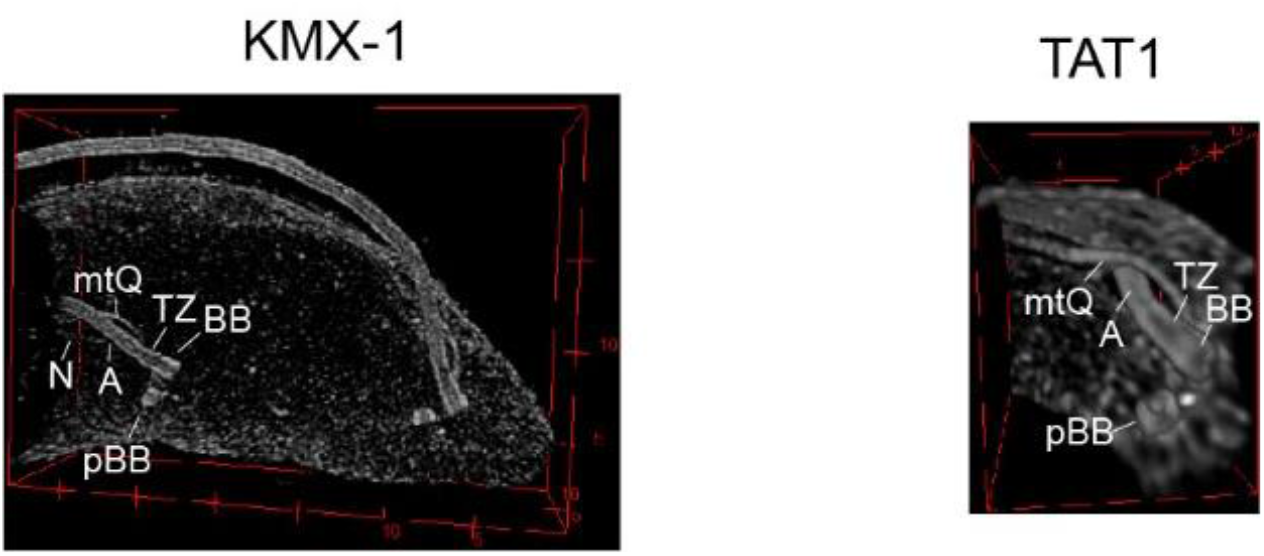
3D reconstruction of the BB area of an expanded procyclic *T. brucei* cell stained with KMX-1 (left; also shown in Suppl. fig. 1A) or with TAT1 (right; also shown in Suppl. fig. 1B). Labelled are structures associated with the old flagellum. A - axoneme; BB - basal body; pBB - pro-basal body; mtQ - microtubule quartet; TZ - transition zone; N - neck microtubule.

**Supplementary figure 7.**
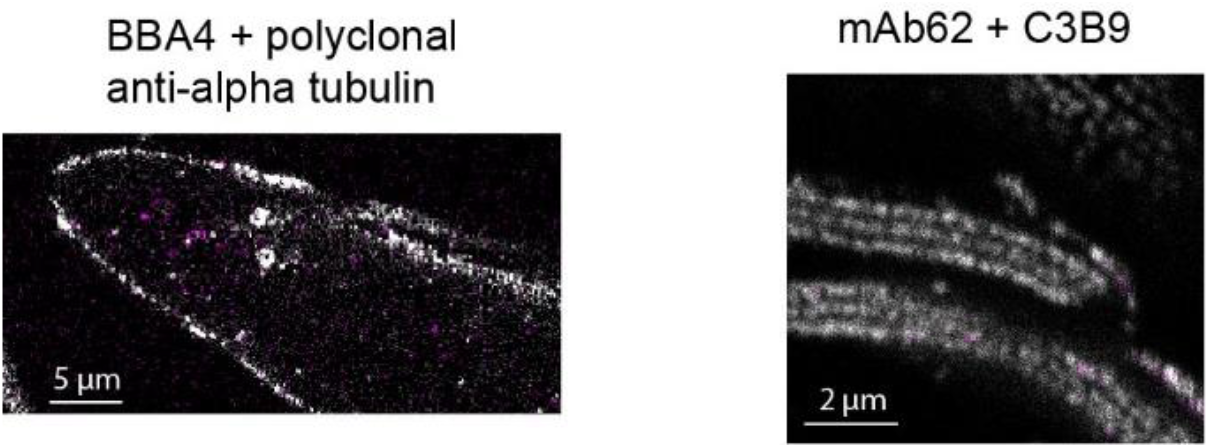
Individual confocal planes of expanded procyclic *T. brucei* cells showing a lack of specific staining (magenta) of the BB and pBB by the antibody BBA4 (left) and of the flagella connector at the tip of the new flagellum axoneme by mAb62 (right). Signals of anti-tubulin antibodies shown in grey.

